# Astrocytes control motor neuronal mitochondrial axonal transport deficits in C9ORF72 ALS

**DOI:** 10.1101/2025.06.10.658820

**Authors:** Maria Stavrou, Áine Heffernan, Roderick N. Carter, Owen Dando, Zoeb Jiwaji, Karen Burr, Jyoti Nanda, Justyna Cholewa-Waclaw, David Story, Giles E. Hardingham, Siddharthan Chandran, Bhuvaneish T. Selvaraj

**Affiliations:** UK Dementia Research Institute at University of Edinburgh, University of Edinburgh, Edinburgh BioQuarter, Chancellor’s Building, 49 Little France Crescent, Edinburgh, EH16 4SB, UK; Centre for Clinical Brain Sciences, University of Edinburgh, Edinburgh, UK; Anne Rowling Regenerative Neurology Clinic, University of Edinburgh, Edinburgh, UK; Euan MacDonald Centre for MND Research, University of Edinburgh, Edinburgh, UK; Centre for Discovery Brain Sciences, University of Edinburgh, Edinburgh, UK; Institute for Regeneration and Repair, University of Edinburgh, Edinburgh, UK

## Abstract

Disrupted axonal transport and astrocyte dysfunction are implicated in amyotrophic lateral sclerosis (ALS). Here, we show these are intrinsically linked: human induced pluripotent stem cell-derived astrocytes carrying the C9ORF72 mutation disrupt axonal transport and mitochondrial function in motor neurons (MNs). Conversely, either isogenic gene-corrected astrocytes or selectively boosting C9ORF72-astrocyte mitochondrial bioenergetics rescue axonal transport deficits in C9ORF72-MNs. Thus, astrocytes have a dominant effect on axonal mitochondrial transport in the context of ALS.

## Main text

Amyotrophic lateral sclerosis (ALS) is a progressive, incurable, and fatal neurodegenerative condition. Mutation in the C9ORF72 gene is the commonest cause of familial-ALS and accounts for c.10% of sporadic cases. Axonal transport of cargo such as mitochondria is essential for neuronal function and survival (1,2). Moreover, genetic and *in vivo* evidence suggests a potential causative link between dysregulated axonal transport and ALS (3). Specifically, abnormal mitochondria function and motility are described in ALS including human autopsy and cellular studies of C9ORF72 MNs (4,5). Dysfunctional mitochondrial bioenergetics in C9ORF72 MNs have been shown to be a key mediator in disrupting axonal transport, whereby enhancing MN mitochondrial function was sufficient for recovery of axonal transport in C9ORF72 MNs (6). Concurrently, astrocytes are key regulators of neuronal metabolism and their role in mediating non-cell autonomous neurodegeneration has been established by multiple lines of experimental and pathological evidence (7,8). Several mechanisms have been described including reduced expression of the glutamate uptake transporter EAAT2 causing downregulation of the glutamate-glutamine cycling and hyperexcitability (9), defects in astrocyte-neuronal lactate shuttling (7) and disrupted membrane transport of mitochondrial-specific energy substrates with consequential loss of metabolic flexibility in C9ORF72 astrocytes (10). However, little is known about astrocyte involvement in controlling axonal transport, with studies to date, largely exploring the intraneuronal consequences of genetic mutations on axonal trafficking. Against this background, we used human stem cell models to address if and how astrocytes regulate axonal mitochondrial transport in the context of C9-ALS. We uncovered a hitherto unreported mechanism of the modulation of axonal transport through astrocytes and found surprising evidence that boosting astrocytic mitochondrial metabolism is sufficient to revert axonal transport deficits in C9ORF72 MNs.

To evaluate cell-autonomous and astrocyte-derived non-cell autonomous effects of the C9ORF72 mutation on mitochondrial axonal transport in MNs, we performed live imaging of mitochondrial transport in MNs, both in isolation and in co-culture with astrocytes.We used three patient-derived human induced pluripotent stem cell (hiPSC) lines carrying the C9ORF72 mutation (C9) and their isogenic gene-corrected controls (C9Δ) generated via CRISPR/Cas9 (11), to produce highly enriched spinal cord-patterned astrocytes expressing >90% GFAP and S100B **(Suppl Fig 1)** and MNs, as previously described (11,12). As we published, notwithstanding that these astrocytes exhibit key C9ORF72-related pathological features, such as RNA Foci and di-peptide repeat proteins, they show no difference in their ability to take up glutamate and propagate calcium waves (11). C9 and C9Δ patient-derived MNs (C9-MN or C9Δ-MN) were examined in monocultures or in co-culture with the three pairs of C9 and C9Δ patient-derived astrocytes (C9-A or C9Δ-A). We measured two established features of mitochondrial axonal transport: the proportion and average velocity of motile mitochondria. First, we selectively labelled control MNs (C9Δ-MNs or human embryonic stem cell derived *HB9::EGFP* [hESC *HB9::EGFP* MNs]) with mito-dsRed2 for measuring axonal mitochondrial transport and then co-cultured them with C9-A. Co-culturing control MNs with mutant astrocytes resulted in significant reductions in both the percentage and speed of motile mitochondria in MNs **(Fig 1a, b, c, e; Suppl Fig 2a; Suppl video 1)**, compared to monocultures of control MNs **(Suppl video 2)**. Significantly, both mitochondrial transport phenotypes were reversed when control MNs (C9Δ-MNs or hESC *HB9::EGFP* MNs) were co-cultured with control C9Δ-Α **(Fig 1a, b, c, e; Suppl Fig 2a; Suppl video 3)**. We next treated control motor neurons (hESC *HB9::EGFP* MNs) with conditioned medium from both C9 and C9Δ astrocytes. The absence of significant changes in mitochondrial motility **(Suppl Fig 2b)** indicates that direct contact, but not secreted factors from astrocytes, is required for the observed axonal transport deficits. We next examined if co-cultures of mutant MNs with mutant astrocytes would exacerbate axonal transport deficits, noting earlier reports of cell-autonomous axonal transport impairments in C9-MNs (5,6). Indeed, in comparison with isolated C9-MNs **(Suppl video 4)**, we observed a significant decrease in the speed of motile mitochondria when C9-MNs were co-cultured with C9-A; although the percentage of motile mitochondria was not affected **(Fig 1d, f; Suppl video 5)**. Importantly, co-culturing C9-MNs with control C9Δ-Α rescued both axonal transport deficits (% motility and mean speed of mitochondria) to wildtype levels **(Fig 1b, d, f; Suppl video 6)**. These data recapitulate previous findings of cell-autonomous axonal transport deficits of C9ORF72 mutation on MNs and further indicate that in co-culture, the MN axonal transport phenotype is determined by the genotype of the astrocyte rather than that of the neurons.

**Figure 1.**
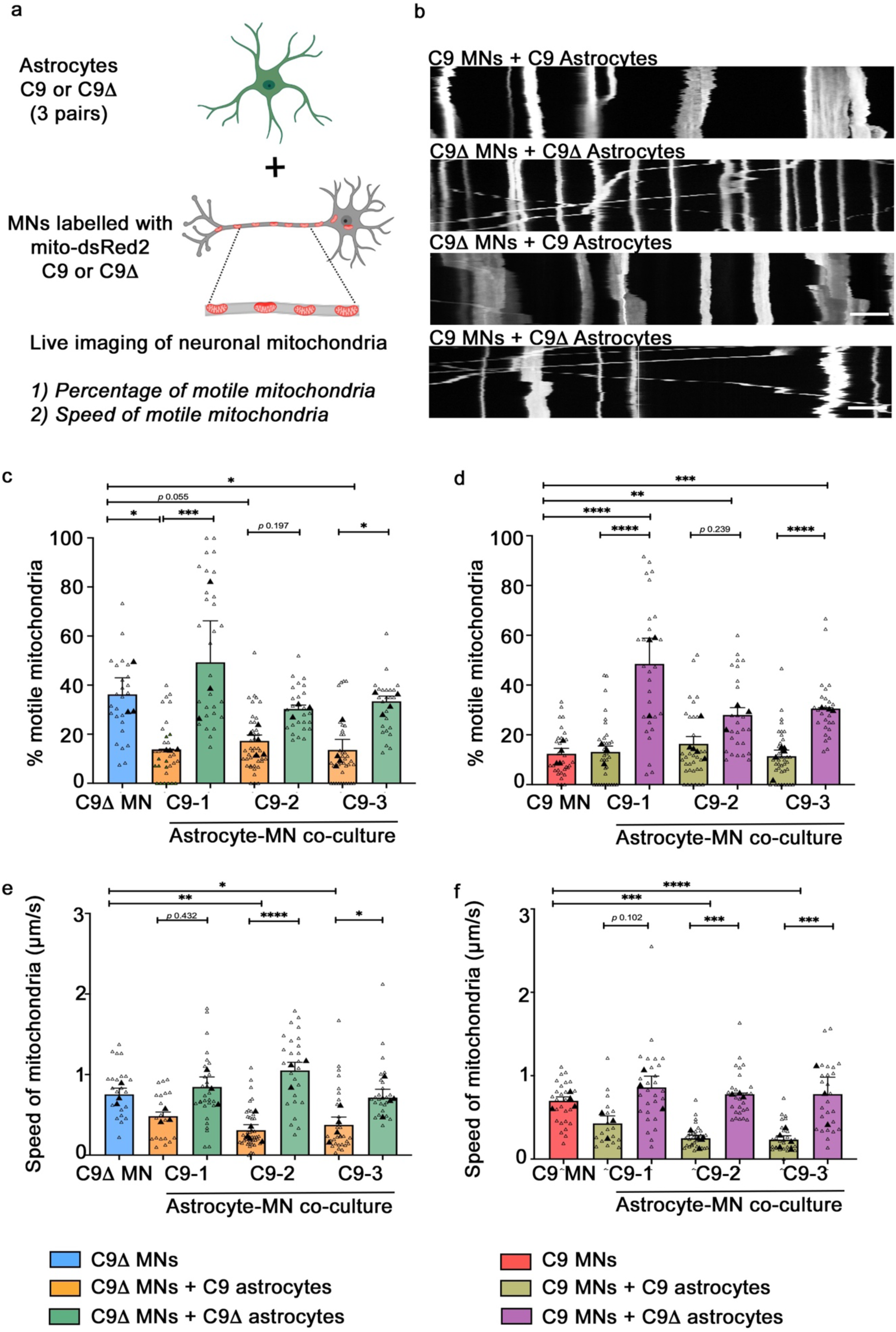
Effects of astrocytes on mitochondrial transport in motor neurons in physical co- cultures. **(a)** Schematic diagram of MN and astrocyte physical co-culture platform to assess mitochondrial transport in MNs. **(b)** Kymographs depicting mitochondrial transport in motor neurons upon co-culture with astrocytes over a 100 µm stretch of the proximal axon of MNs; scale bar = 10 μm. **(c)** Quantification of the percentage of motile mitochondria relative to the total number of mitochondria labelled with mitoDsRed2 over a 100 µm stretch of the proximal axon of C9Δ MNs either in isolation or in co-culture with C9 astrocytes and their gene- corrected paired controls across the three independent pairs. **(d)** Quantification of the percentage of motile mitochondria relative to the total number of mitochondria labelled with mitoDsRed2 over a 100 µm stretch of the proximal axon of C9 MNs either in isolation or in co- culture with C9 astrocytes and their gene-corrected paired controls across the three independent pairs. **(e)** Measurement of the mean speed of motile mitochondria over a 100 µm stretch of the proximal axon of C9Δ MNs either in isolation or in co-culture with C9 astrocytes and their gene-corrected paired controls across the three independent pairs. **(f)** Measurement of the mean speed of motile mitochondria over a 100 µm stretch of the proximal axon of C9 MNs either in isolation or in co-culture with C9 astrocytes and their gene-corrected paired controls across the three independent pairs. MNs throughout the study were generated from one pair of C9 mutant and isogenic gene-corrected control line. C9-A (1–3) represents mutant astrocytes, C9-MN denotes mutant motor neurons, C9Δ-A (1–3) indicates isogenic gene- corrected astrocytes, and C9Δ-MN refers to isogenic gene-corrected motor neurons. The open triangles represent technical replicates (individual axons) and the filled triangles represent independent biological replicates (mean of technical replicates). Data are represented as the mean of biological replicates ± SEM of the means of biological replicates. p-values (*<0.05; **<0.01; ***<0.001; ****<0.0001) determined by fitting generalised linear mixed models, followed by post-hoc tests with Bonferroni correction.

Given the established role of astrocytes in regulating brain energy homeostasis (13,14), we subsequently examined their direct impact on mitochondrial bioenergetics in MNs using the Seahorse assay in co-cultures. Our comparison of mitochondrial oxygen consumption rates (OCR) revealed that MN monocultures and MNs co-cultured with astrocytes exhibited nearly 8-fold higher OCR than astrocytes alone **(Suppl Fig 3a)**, implicating that any significant changes in OCR values in co-cultures, are primarily driven by MNs. This conclusion is further supported by MitoTracker Red CMXROS assays (4,15), which showed no difference in astrocytic steady-state mitochondrial membrane potential (MMP) between mono- and co-culture conditions **(Suppl Fig 3b).** We corroborated previous findings indicating impairments in both basal and maximal mitochondrial respiration in monocultures of C9-MNs with an observed reversal in C9-Δ MNs **(Suppl Fig 4b-c)** (6). Our results showed that co-culturing MNs with astrocytes, irrespective of the astrocyte genotype, enhanced basal and maximal respiration compared to MN monocultures, despite comparable neuronal densities **(Suppl Fig 4a-c)**. Crucially, we found reductions in basal and maximal respiration when either C9-MNs or C9Δ-MNs were co-cultured with C9-A in comparison to co-culture with C9Δ-A **(Fig 2a-b)**. This observation indicates that the astrocyte genotype modulates neuronal mitochondrial bioenergetic function.

**Figure 2.**
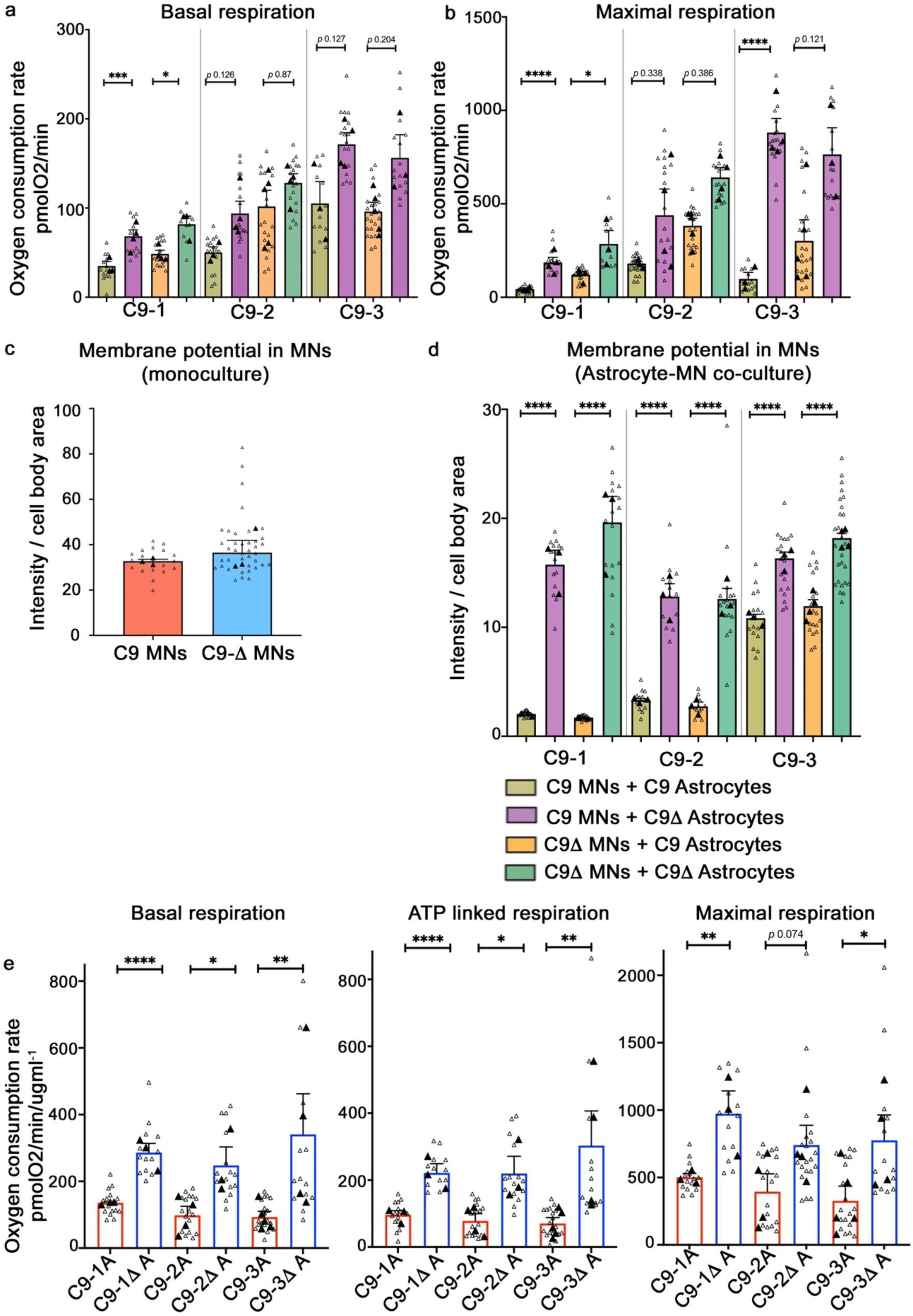
C9 astrocytes impair mitochondrial bioenergetics in MNs. **(a) & (b)** Quantification of the OCR as measured by the Seahorse Analyzer denoting basal and maximal FCCP-uncoupled mitochondrial respiration, involving co-cultures of astrocytes and MNs derived from patient-specific hiPSC lines. The astrocytes were obtained from three C9 hiPSC lines and their corresponding isogenic gene-corrected counterparts created via CRISPR/Cas9. MNs were generated from one pair of C9 mutant and isogenic gene-corrected control lines. The open triangles represent technical replicates and the filled triangles represent independent biological replicates (mean of technical replicates). Data are represented as the mean of biological replicates ± SEM of the means of biological replicates. p-values (*<0.05; **<0.01; ***<0.001; ****<0.0001) determined by fitting generalised linear mixed models, followed by post-hoc tests with Bonferroni correction. **(c)** Quantification of the mean fluorescence intensity of MitoTracker Red CMXROS in hiPSC-derived C9 and C9Δ MNs. The open triangles represent technical replicates and the filled triangles represent independent biological replicates (mean of technical replicates). In total, over 20 MNs per well were analysed to evaluate intensity per cell body area (automated analysis). Data are represented as the mean of biological replicates ± SEM of the means of biological replicates**. (d)** Quantification of the mean fluorescence intensity of MitoTracker Red CMXROS in hiPSC- derived C9 and C9Δ MNs co-cultured with hiPSC-derived C9 A and C9A astrocytes (from the three-independent patient-derived C9ORF72 lines). The open triangles represent technical replicates and the filled triangles represent independent biological replicates (mean of technical replicates). In total, over 20 MNs per well were analysed to evaluate intensity per cell body area (automated analysis). Data are represented as the mean of biological replicates ± SEM of the means of biological replicates. p-values (*<0.05; **<0.01; ***<0.001; ****<0.0001) determined by fitting generalised linear mixed models, followed by post-hoc tests with Bonferroni correction. **(e)** C9 astrocytes, when compared to their corresponding gene-edited controls, display defective bioenergetics (diminished capacity to produce ATP). OCR as measured by the Seahorse Analyzer, normalised to the amount of total protein, denoting basal, ATP-linked, and maximal FCCP-uncoupled mitochondrial respiration for C9 astrocytes and gene-corrected paired controls across the three independent pairs. The open triangles represent technical replicates, and the filled triangles represent independent biological replicates (mean of technical replicates). Data are represented as the mean of biological replicates ± SEM of the means of biological replicates. p-values (*<0.05; **<0.01; ***<0.001; ****<0.0001) determined by fitting generalised linear mixed models, followed by post-hoc tests with Bonferroni correction.

To confirm the effect of astrocyte co-culture on neuronal mitochondrial function we next assessed steady-state MMP in MNs with MitoTracker Red CMXRos. MNs were labelled using lentiviruses expressing GFP or identified using immunostaining with neuron-specific marker SMI312 post-fixation. Quantification of the mean fluorescence intensity (normalised to ROI) of MitoTracker Red CMXROS in MNs (monocultures) showed reduced MMP in the C9-MN line compared to its paired gene-corrected control **(Fig 2c)**. In co-culture with C9-A, the MMP of C9-MN or C9Δ-MNs was significantly reduced. Importantly, this deficit was reversed in co-culture with their corresponding C9Δ-A **(Fig 2d)**. Taken together, these findings demonstrate for the first time, that astrocytes harbouring C9ORF72 mutation, upon co-culture, directly mediate perturbations in the mitochondrial bioenergetics of control and mutant MNs.

Having established that astrocytes directly modulate MN mitochondrial bioenergetics, we next examined the impact of C9ORF72 mutation in isolated astrocytes on mitochondrial bioenergetics using the Seahorse assay. This revealed that C9-A compared to their paired gene-corrected controls, displayed significantly reduced basal, ATP-linked, and maximal respiration **(Fig 2e)**. Therefore, we hypothesised that bioenergetic deficits in C9-A are responsible for their non-cell autonomous effects on axonal transport in MNs. To test this hypothesis, we first overexpressed transcriptional co-regulator peroxisome proliferator-activated receptor gamma co-activator 1-alpha (PGC1α) – a central inducer of mitochondrial biogenesis and metabolism – in C9-A using lentivirus (16,17), which significantly increased OCR in isolated C9-astrocyte cultures **(Suppl Fig 5).** We then co-cultured transduced C9-A with either C9-MNs or C9Δ-MNs and found that the deficits in mitochondrial motility and speed were rescued **(Fig 3a-e; Suppl video 7; Suppl video 8)**. In summary, we uncover that in C9ORF72-ALS, astrocyte-specific rescue of mitochondrial bioenergetics with PGC1α is sufficient to reverse axonal transport deficits in C9ORF72 MNs, revealing non-cell autonomous metabolic regulation of astrocytes on axonal transport.

**Figure 3.**
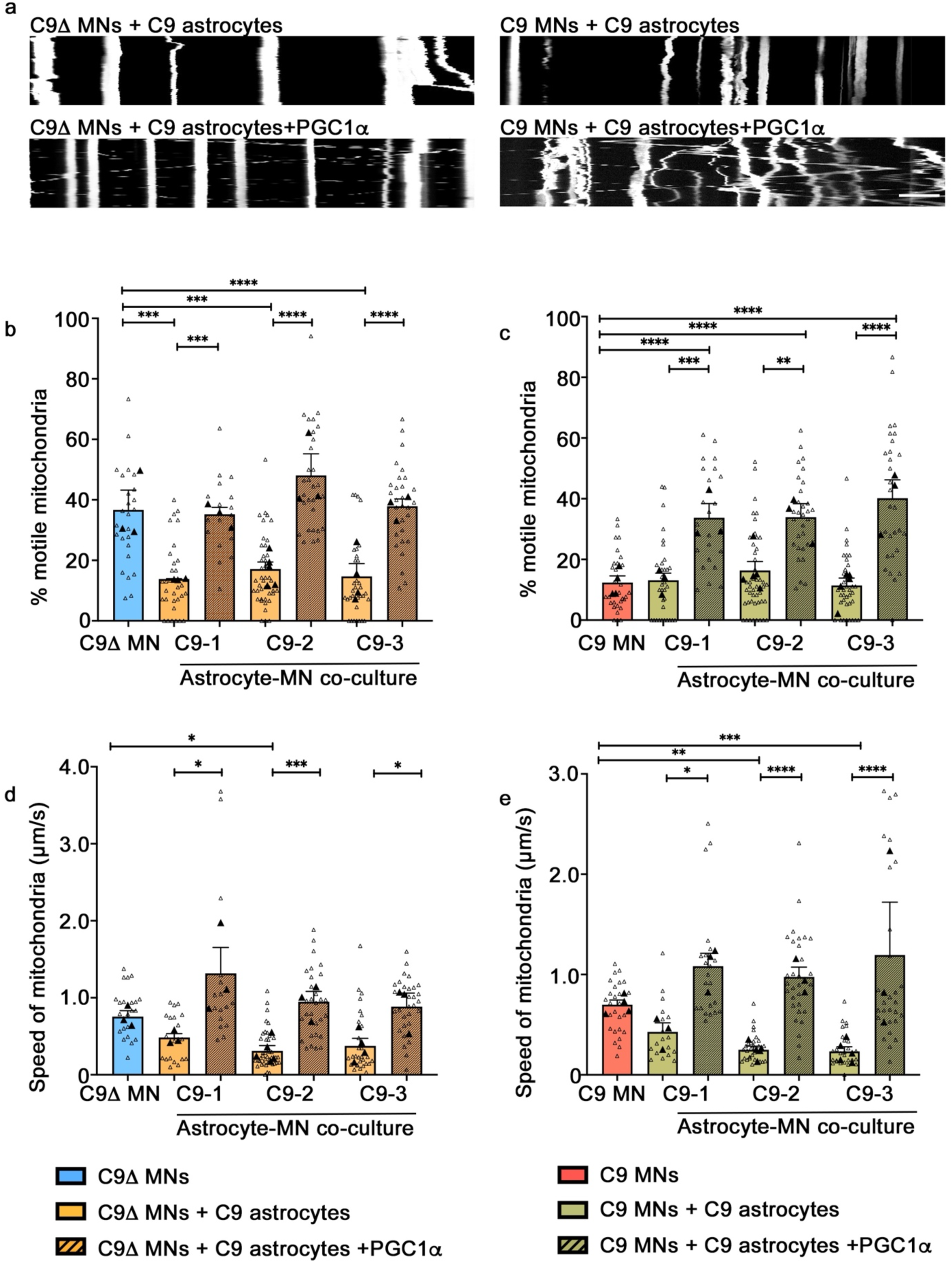
Boosting astrocyte metabolism with selective PGC1α pathway manipulation in C9 astrocytes rescues the neuronal mitochondrial transport deficit. **(a)** Kymographs depicting mitochondrial transport in motor neurons upon co-culture with C9-A and C9-A overexpressing PGC1a; scale bar = 10 μm. **(b)** Quantification of the percentage of motile mitochondria relative to the total number of mitochondria labelled with mitoDsRed2 over a 100 µm stretch of the proximal axon of C9Δ MNs either in isolation or in co-culture with either C9-A with or without PGC1α overexpression from the three independent patient lines. **(c)** Quantification of the percentage of motile mitochondria relative to the total number of mitochondria labelled with mitoDsRed2 over a 100 µm stretch of the proximal axon of C9 MNs either in isolation or in co- culture with either C9-A with or without PGC1α overexpression from the three independent patient lines. **(d)** Measurement of the mean speed of motile mitochondria over a 100 µm stretch of the proximal axon of C9Δ MNs either in isolation or in co-culture with either C9-A with or without PGC1α overexpression from the three independent patient lines. **(e)** Measurement of the mean speed of motile mitochondria over a 100 µm stretch of the proximal axon of C9 MNs either in isolation or in co-culture with either C9-A with or without PGC1α overexpression from the three independent patient lines. MNs throughout this study were generated from one pair of C9 mutant and isogenic gene-corrected control line. The open triangles represent technical replicates (individual axons) and the filled triangles represent independent biological replicates (mean of technical replicates). Data are represented as the mean of biological replicates ± SEM of the means of biological replicates. p-values (*<0.05; **<0.01; ***<0.001; ****<0.0001) determined by fitting generalised linear mixed models, followed by post-hoc tests with Bonferroni correction.

## Discussion

This study reveals a novel non-cell autonomous mechanism regulating axonal transport, whereby C9ORF72 mutant astrocytes impair the proportion and average velocity of motile mitochondria in MNs. This contrasts with the cell-autonomous deficits in C9 MNs, where speed of motile mitochondria is unaffected (9,10), suggesting distinct astrocyte-mediated mechanisms. While multiple factors influence mitochondrial velocity, including localisation of the cargo and the directionality of the transport (18,19), our data reveal a distinct role for astrocytes.

Importantly, C9 astrocytes exhibit reduced glycolytic capacity, lactate production **(Suppl Figure 6)**, and mitochondrial respiration **(Fig 2e)** indicating bioenergetic deficiency. RNA sequencing did not show genotype-driven astrocytic reactivity **(Suppl Figure 7)**, excluding a gain-of-toxic-function mechanism. This suggests a loss of astrocytic physiological function and impaired metabolic support for neurons, potentially due to: (a) a defective lactate shuttle compromising axonal ATP generation; and/or (b) mitochondrial dysfunction, disrupting essential astrocytic functions, including glutamate regulation, calcium signaling, transmitophagy, mitochondrial transfer and fatty acid metabolism (20–22). The precise mechanisms underlying this astrocyte-mediated axonal mitochondrial transport dysfunction require further investigation. Beyond mitochondrial motility, future studies should explore astrocytic regulation of other ALS-relevant organelle transport (e.g., peroxisomes, lysosomes, RNP complexes). The broader relevance of our findings to other ALS mutations and sporadic ALS also requires investigation. Our findings, showing that upregulating PGC1α in C9ORF72 astrocytes improves axonal transport, support its therapeutic potential in ALS. This aligns with studies demonstrating that PGC1α activation in muscles of SOD1 mice maintained mitochondrial biogenesis and activity, accompanied by delayed muscle atrophy and improved muscle endurance even at late stages of disease (23).

In summary, we report a novel role of astrocytes in regulating axonal transport through mitochondrial bioenergetics, thus, providing evidence for astrocyte metabolism as a potential ALS therapeutic target.

## Methods

### Generation of hiPSC lines and isogenic gene-corrected control hiPSC lines

Dermal fibroblasts from three C9ORF72 ALS/FTD patients (C9-1, C9-2, C9-3) harbouring the G4C2 repeat expansion in the C9ORF72 gene were acquired and reprogrammed under full Ethical/Institutional Review Board approval at the University of Edinburgh. Details of the C9 hiPSC lines including clinical meta-data of the patients are listed in *Mehta et al.,2021* (6). Gene-edited controls were generated via CRISPR/Cas9 technology, as previously described by our group (6,11,12). Over the course of this study, karyotypes of all hiPSC lines were examined using the conventional Giemsa banding chromosomal analysis, conducted by The Doctors Laboratory Ltd, London. Exclusion of mycoplasma contamination was ascertained through monthly testing of hiPSC culture supernatants using the Venor^®^GeM Classic detection kit (Minerva Biolabs GmbH). hiPSC lines were cultured with Essential 8 medium and passaged using Dispase/Collagenase.

### hiPSC motor neuron generation

Spinal MN differentiation from parental and gene-edited hiPSCs (one C9-mutant/gene-corrected MN pair was used throughout the study) was performed using an established protocol with minor modifications, yielding a highly enriched and electrophysiologically mature neuronal culture, devoid of glia, with approximately 60% of cells being positive for islet-1 and islet-2 homeobox MN markers 1-week post-plating (6). hiPSCs were dissociated into single cells using 1X Accutase® (Sigma-Aldrich) and neuralised as a suspension culture utilizing dual-SMAD inhibition with SB-431542 (20 μM; Tocris Bio-Techne), LDN-193189 (0.1 μM; Selleckchem), and potentiation with the Wnt-agonist, CHIR- 99021 (3 μM; Tocris Bio-Techne) in N2/B27 medium, which consists of 0.5X Neurobasal™ (Gibco Thermo Fisher Scientific), 0.5X Advanced DMEM/F12 (Gibco Thermo Fisher Scientific), 1X Antibiotic–Antimycotic (Gibco Thermo Fisher Scientific), 1X GlutaMAX™ (Gibco Thermo Fisher Scientific), 100 μM beta-mercaptoethanol (Gibco Thermo Fisher Scientific), 1X B-27™ supplement (Gibco Thermo Fisher Scientific), 1X N-2 supplement (Gibco Thermo Fisher Scientific), and 10 μM l-ascorbic acid (Sigma-Aldrich). On day 2, neural spheres were simultaneously patterned to spinal cord identity by treatment with retinoic acid (RA, 0.1 μM; Sigma-Aldrich) and smoothened agonist (SAG, 0.5 μM; Sigma-Aldrich), promoting caudalisation and ventralisation, respectively, along with SB-431542, LDN-193189, and CHIR-99021 in N2/B27 medium for an additional 5 days. On day 7, spheres were maintained in culture with RA and SAG, supplemented with recombinant human brain-derived neurotrophic factor (BDNF, 10 ng/ml; R&D Systems Bio-Techne) in N2/B27 medium to generate MN progenitors. From day 9, MN progenitors were cultured in the day 7 medium with the addition of DAPT (10 μM; Tocris Bio-Techne), an inhibitor of Notch signaling, for an additional 5 to 7 days. At days 14 to 16, MN spheres were dissociated using 0.05% Trypsin– EDTA (Gibco Thermo Fisher Scientific), and cells were plated as a monolayer onto dishes coated with laminin from Engelbreth-Holm-Swarm murine sarcoma basement membrane (5 μg/mL; Sigma-Aldrich), fibronectin human plasma (10 μg/ml; Sigma-Aldrich), and Matrigel® (1:20), which had been pre-treated with poly-l-ornithine bromide (100 µg/ml; Sigma-Aldrich). On day 0, the cells were cultured in motor neuron neurotrophic factor (MN-NF) medium, which contains 1X Neurobasal, 1X Antibiotic–Antimycotic, 1X GlutaMAX™, 1X MEM Non-Essential Amino Acids solution (Gibco Thermo Fisher Scientific), 100 μM beta-mercaptoethanol, 1X B- 27™ supplement, 1X N-2 supplement, RA (1 μM), ascorbic acid (2.5 μM), BDNF (10 ng/ml), recombinant human glial-derived neurotrophic factor (GDNF, 10 ng/ml; R&D Systems Bio-Techne), recombinant human ciliary neurotrophic factor (CNTF, 10 ng/ml; R&D Systems Bio-Techne), and animal-free recombinant human insulin-like growth factor-1 (IGF-1, 10 ng/ml; PeproTech), along with 25 μM uridine/5-fluoro-2′-deoxyuridine (U/FDU). From day 2 onwards, the medium was regularly changed every 2 to 3 days and supplemented with uridine/5-fluoro- 2′-deoxyuridine (1 μM; Sigma-Aldrich) for at least 1-week post-plating, to eliminate residual proliferating cells. For MNs, independent biological replicates/plate downs refer to distinct neuralisation and differentiation steps originating from the hiPSC state.

### hiPSC astrocyte generation

Astrocytes were generated from hiPSCs using established lab protocols and were characterised through expression of astrocyte-specific markers and implementation of functional assays (glutamate uptake assay and calcium imaging) as described earlier publications (11). Specifically, hiPSCs were neuralised and subsequently converted into spheres following the established protocols described in the previous section. The spheres were then cultured in MN maturation medium for a duration of 2 to 4 weeks. Following this maturation phase, the spheres were chopped and cultured in NSCR EL20 medium for an additional 4 to 6 weeks to induce astrogliogenesis. At the conclusion of this conversion phase, the medium was switched to NSCR EF20 medium to support the proliferation of astrocyte progenitor cells (APCs) within the spheres. These astrospheres were mechanically chopped every 4 weeks to ensure good viability and maintained long-term in EF20 medium (>4 chops). The resulting astrospheres were dissociated into single cells using the Papain Dissociation System (Worthington Biochemical) and plated onto 6-well plates coated with Matrigel (BD Biosciences, diluted 1:80) at a density of 7.5 × 10^5^ cells per well.

These cells were subsequently differentiated into astrocytes by transitioning the medium to AstroMed CNTF medium for a period of 2 weeks. Throughout the astrocyte generation process, the media were changed every 2 to 3 days to ensure optimal growth conditions. The NSCR EF20 medium comprised Advanced DMEM/F12 (Invitrogen), 1% N2 supplement (Invitrogen), 0.1% B-27 supplement (Invitrogen), 1% penicillin/streptomycin (Invitrogen), 1% GlutaMAX solution (Invitrogen), 20 ng/ml fibroblast growth factor 2 (FGF-2; PeproTech), and 20 ng/ml epidermal growth factor (EGF; R&D Systems). The AstroMed CNTF medium consisted of Neurobasal medium (Invitrogen), 0.2% B-27 supplement (Invitrogen), 1% non-essential amino acids (NEAA; Invitrogen), 1% penicillin/streptomycin (Invitrogen), 1% GlutaMAX (Invitrogen), and 10 ng/ml CNTF (R&D Systems).

We used 3 patient-derived iPSC astrocyte lines carrying the C9ORF72 mutation (C9), along with their isogenic gene-corrected control with the C9ORF72 mutation removed (C9Δ) using CRISPR/Cas9. For each hiPSC line, astrocyte progenitor cells (APCs) were derived and banked from two separate dissociations of astrospheres. For each biological experiment/plate down, APCs were thawed out and differentiated independently into astrocytes over two weeks.

### Co-cultures of MNs and astrocytes

Astrocyte progenitor cells were dissociated into single cells and plated in Astrocyte differentiation medium (AstroMed CNTF medium) to differentiate into mature astrocytes over 2 weeks, as described above. At 2 weeks, MN spheres (day 16) were dissociated into single cells using 0.05% Trypsin/EDTA and plated on astrocyte monolayers in MN-NF medium (1X Neurobasal, 1X Antibiotic–Antimycotic, 1X GlutaMAX™, 1X MEM Non-Essential Amino Acids solution [Gibco™ Thermo Fisher Scientific], 100 μM beta-mercaptoethanol, 1X B-27™ supplement, 1X N-2 supplement, RA [1 μM], ascorbic acid [2.5 μM], recombinant human BDNF [10ng/ml,248-BD/CF, R & D Systems^®^ Bio-Techne], recombinant human GDNF [10 ng/ml; 212-GD, R & D Systems^®^ Bio-Techne], recombinant CNTF [10 ng/ml; 257-NT, R & D Systems^®^ Bio-Techne], and animal-free recombinant human IGF-1 [10 ng/ml; AF-100–11, PeproTech^®^]).

Astrocyte-MN co-cultures and MN monocultures were maintained for 3 weeks in MN-NF medium prior to performing experiments. MN-NF medium was changed every 2–3 days and supplemented with U/FDU (1 μM, U3003 and F0503, Sigma-Aldrich^®^) for at least 1-week after plate-down to remove residual proliferating cells.

**Table 1:**
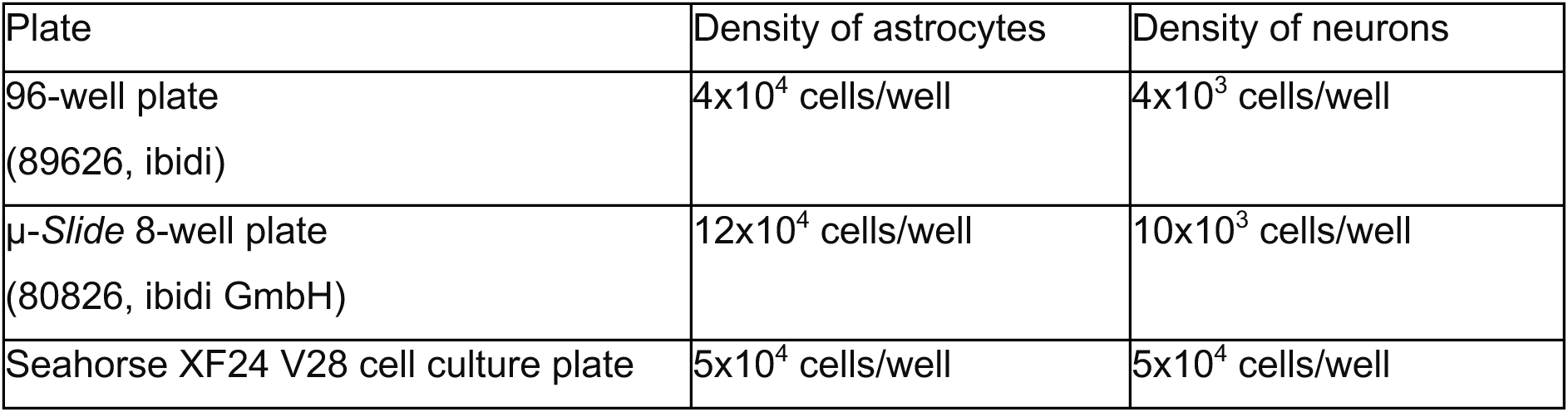
Density of neurons and astrocytes in physical co-cultures.

### Axonal Transport Studies

Astrocyte-MN co-cultures were generated on μ-*Slide* 8-well plates as described above. MN monocultures were also produced by plating MNs as a monolayer on µ-slide 8-well ibidi dishes as explained in prior work (6,24). At plate-down, the MNs were sparsely transduced with lentivirus expressing mitoDsRed2 as follows:

a. MN Monocultures: Multiplicity of infection (MOI) of 0.5-1, adjusted to obtain 1-2 labelled cells per field of view, as described previously (6).
b. Astrocyte-MN co-cultures: mitoDsRed2 lentivirus was added at MOI 10 to approximately 10,000 MNs (per well; on μ-*Slide* 8-well plates) and MNs were selectively transduced prior to being transferred onto fully differentiated astrocytes.

Live imaging of mitochondrial axonal transport was undertaken at 21 days post MN plate- down at 63 X magnification (Plan-Apochrat 1.40 NA oil DIC M27 objective, Carl Zeiss) using an Axio Observer Z1 inverted motorised microscope (Carl Zeiss). The microscope was equipped with a Cy3 FL filter-set (Carl Zeiss), Zen 2011 z-stack, time lapse and Definite Focus modules (Carl Zeiss), and an S1 Environmental System (Carl Zeiss) incubation chamber for temperature control at 37 °C and 5% CO2 flow. Time-lapse imaging of mitochondria in axons was conducted as previously described (6,24). For all conditions, measurements were acquired from the proximal portion of the axon and the axonal segments selected for recording, were at least 100 μm from the cell soma, to avoid the axon hillock – the part of the axon at the juncture with the cell body. Images were acquired at a capture rate of 0.2 Hz for 5 minutes, covering a ∼ 100 µm stretch of axon and a small z-stack, if required. Motile mitochondria were defined as those travelling at or above the cut-off speed of 0.1 µm/s (5). Time lapse images were used to generate videos, exported from the Zeiss software and further analysis was implemented in Fiji. Kymographs were generated and analysed using KymoToolBox ImageJ plugin (25) to determine the numbers of static (≤ 0.1 µm/s) *versus* bidirectionally motile mitochondria (either towards or away from the soma). A minimum of 4 axons (n) were imaged per condition per biological replicate (N, where N=3). The percentage of motile mitochondria (labelled with mitoDsRed2) relative to the total number of mitochondria was quantified in a 100 µm stretch of axon in *C9orf72* and gene-edited MNs. For calculating the mean speed of motile mitochondria in each segment measured, only mitochondria with a speed higher than 0.1 μm per second over the recorded section of time (5 minutes) were included in the analysis. The average speed of those mitochondria per axon was taken forward.

### Metabolic profiling of astrocytes & neurons (seahorse assay)

The following cell densities were used for the seahorse assay:

a. Astrocyte monocultures: APCs were plated on V7 Seahorse plates (100777-004, Agilent) at a density of 5x10^4^ cells per well to ascertain a highly confluent cellular monolayer. They were cultured in CNTF medium for differentiation into astrocytes over 2 weeks.
b. MN monocultures: MNs were plated on poly-L-ornithine bromide (100 µg/ml; P3655, Sigma-Aldrich^®^)-treated V28 Seahorse plates (100882-004, Agilent). The plates were subsequently coated overnight as described in *Mehta et al., 2021* (6). Neuronal density was at 50,000 cells per well.
c. Astrocyte-MN co-cultures: APCs were dissociated into single cells and plated at a density of 5x10^4^ cells per well on V28 Seahorse plates in CNTF to differentiate progenitors into mature astrocytes over 2 weeks. At 2 weeks, MN spheres (day 16) were dissociated into single cells using 0.05% Trypsin/EDTA and plated on astrocyte monolayers in MN-NF medium at a density of 5x10^4^ cells per well.

The standard Agilent Seahorse XF Cell Mito Stress protocol was modified depending on the condition and the experiments were conducted at 21 days post MN plate down using a Seahorse XFe24 Analyzer (Agilent).

### Mitochondrial Function

The Seahorse reader can measure OCR, an excellent read-out for mitochondrial respiratory activity and the current choice experiment to examine underlying mitochondrial dysfunction. Culture medium was exchanged for low-buffering capacity Seahorse XF Base Medium (102,353, Agilent) supplemented with 1X Glutamax™, 10 mM glucose (Sigma-Aldrich^®^) and 2 mM pyruvate (Sigma-Aldrich^®^), adjusted to pH 7.35 ± 0.5 at 37 °C with 1M sodium hydroxide. The cells were equilibrated for 30 minutes at 37 °C with no CO2, prior to being inserted in the machine. OCR was measured basally and after sequential addition of specific mitochondrial inhibitors as illustrated in the table below. FCCP dose was optimised for astrocyte and neuronal monocultures (tested doses for neurons – 0.125 μM, 0.25 μM, 0.5 μM; tested doses for astrocytes – 0.125 μM, 0.25 μM, 0.5 μM, 1 μM) and the concentration adopted that maximised the OCR response.

**Table 2a:** Drugs targeting the mitochondrial bioenergetic machinery and their concentrations.

| Drug | Target | Astrocyte monocultures | MN monocultures | Astrocyte-neuronal cocultures |
| --- | --- | --- | --- | --- |
| Oligomycin (Sigma-Aldrich<sup>®</sup>) | F<sub>1</sub>F<sub>0</sub> ATP synthase inhibition | 1.5 μM | 3 μM | 3 μM |
| Carbonyl cyanide 4-(trifluoromethoxy)phenylhydrazone (FCCP; Cayman Chemical, Cambridge Bioscience) | Increases proton leak; uncouples mitochondria | 1 μM | 0.5 μM | 0.5 μM |
| Antimycin/rotenone (Sigma-Aldrich<sup>®</sup>) | Complex III/Complex I inhibitors | 0.5 μM | 3 μM | 3 μM |
| 2-deoxy-D-glucose (D6134, Sigma-Aldrich<sup>®</sup>) | Terminates glycolysis | 100 mM | 100 mM | 100 mM |

The protocol involved a 3-minute mixing, 2-minute wait, 3-minute measure cycle (total duration of each cycle was 8 min). Three measurements/cycle were obtained basally (before drug injections; total duration of 24 min) and three measurements were taken after addition of each drug (first: oligomycin; second: FCCP; third: antimycin/rotenone; fourth: 2-deoxy-D-glucose).

### Glycolytic Rate Measurement

A modified version of the Seahorse Glycolytic Stress test protocol was used as previously described. hiPS-derived astrocyte monocultures, differentiated and plated as above, were washed and incubated in Seahorse Assay Medium supplemented with 1 mM glutamine and with zero glucose for 30 minutes at 37 °C with no CO2. Extracellular acidification rate (ECAR) was measured using the Seahorse pH probe basally and after sequential addition of specific metabolic modifiers at the doses in Table 2b (first: Glucose; second: Oligomycin; third: 2-deoxy-D-glucose), with intermittent mixing and measurement cycles as described above.

**Table 2b:** Drugs used within the glycolytic stress test protocol.

| Drug | Target | Final Concentration |
| --- | --- | --- |
| Glucose (Sigma-Aldrich) | Substrate for glycolysis | 10 mM |
| Oligomycin (Sigma-Aldrich) | ATP synthetase inhibitor | 1.5 μM |
| 2-deoxy-D-glucose (Sigma-Aldrich) | Terminates glycolysis | 100 mM |

Five individual wells were used per condition in each plate, with experiments repeated in three independent plate-downs from different MN differentiations (*N* = 3). In astrocyte and MN monocultures, the measured OCR ([O2]/time) and ECAR were normalised to total protein content using the bicinchoninic acid assay (23225, Pierce™ Thermo Fisher Scientific) (15). For astrocyte-neuronal co-cultures, neuronal counts were estimated through neuron-specific staining [immunocytochemistry with SMI-312 at 1:1000 (837904, BioLegend^®^)]. Data were visualised in, and imported from, Seahorse Wave Desktop software (version 2.6.0.31, Agilent). **Basal respiration,** which represents the net sum of all processes in the cell capable of consuming O2, including mitochondria and other oxidases, was calculated by subtracting non-mitochondrial respiration (i.e., the minimum OCR value after antimycin/rotenone addition) from the OCR value right before oligomycin injection. **Maximal respiration** was calculated by subtracting non-mitochondrial respiration from the highest OCR measurement after FCCP injection. **ATP-linked respiration** was calculated by subtracting the oligomycin rate from baseline cellular OCR. **Glycolytic Capacity** was calculated by subtracting the baseline non- glycolic acidification from the maximum ECAR rate (ECAR differential) achieved following the addition of oligomycin, which stops oxidative phosphorylation and maximising cell glycolysis for energy provision.

### Förster resonance energy transfer (FRET) Metabolic Imaging

hiPSC-derived astrocytes were transfected with plasmids encoding FRET probes for glucose (FLII12Pglu-700uDelta6, #17866) (26), or lactate (Laconic, #44238) (27). This was done 24 h prior to metabolic measurement using Lipofectamine 2000 (Life Technologies) at 0.65 μg of DNA per well and 2.33 μl Lipofectamine 2000 per well for 45 minutes as described previously) (28).

Metabolic imaging was performed with astrocytes being continuously perfused at 37 degrees Celsius with ACSF (composition: 150mM NaCl, 3mM KCl, 10mM HEPES buffer, 0.1mM glycine, 2mM CaCl2, 1mM MgCl2, and 10mM glucose, pH 7.4). Images were captured using a DFC350 FX digital camera as part of a Leica AF6000 LX imaging system. Images were acquired every 10s. All probes were imaged with a standard FRET CFP/YFP filter wheel, with excitation of CFP and measurements taken within the CFP and YFP emission spectra. For glucose measurement, the YFP/CFP ratio of FLII12Pglu-700uDelta6 FRET probe was used to determine intracellular glucose concentrations for individual astrocytes. Levels were measured at baseline before the addition of the glucose-uptake inhibitor cytochalasin B (20uM) after 60s. This results in a reduction in intracellular glucose levels corresponding to the rate of glucose consumption. For lactate measurements, the CFP/YFP ratio of the Laconic FRET probe was used to determine intracellular lactate levels. Levels were measured at baseline in ACSF, before the addition of the MCT inhibitor AR-C155858 (1μM). This results in an increase in concentration corresponding to intracellular lactate production. All FRET ratios were normalised by subtracting baseline and expressed as percentage of maximum. A linear least-squares fitting routine was used to determine the line of best fit and slopes calculated for the portion of the curve corresponding to the rate of consumption or production of substrate.

### Mitochondrial staining for determination of neuronal MMP

Astrocyte-MN co-cultures were generated on 96-well plates as described above. MN monocultures were also produced by plating MNs on 96-well plates at a density of 10,000 cells per well. Neurons were labelled and demarcated from astrocytes either at plate-down with transduction with lentivirus expressing GFP or immunostained after fixation with neuron-specific marker SMI312 at 1:1000 (837904, BioLegend^®^). After 21 days of MN plate down, astrocyte-MN co-cultures and MN monocultures were incubated with 100 nM of potentiometric dye MitoTracker Red CMXRos (Thermo Fisher) for 45 minutes at 37°C (4). This is a membrane-permeant, lipophilic, cationic fluorescent probe that is sequestered in the mitochondrial membrane of living cells and its accumulation is dependent on mitochondrial potential (15). The covalent binding of the dye to the polarised inner mitochondrial membrane permits its retention during washing and fixation of cells and makes it suitable for fluorescence microscopy. The cells were then washed with PBS and fixed in 4% PFA. Fluorescence was visualised using Opera Phenix Plus high-content screening system at 40X magnification (OperaPHX/OPRTCLS Water Immersion Objective 40x: NA 1.1, WD 0.62 mm (field of view: approx. 323 μm x 323 μm). Flatfield correction was automatically performed by Harmony Software during image acquisition. Harmony or Signals Image Artist was used for image analysis.

### Mitochondrial staining for determination of astrocyte MMP

Astrocyte-MN co-cultures and astrocyte monocultures were generated on 96-well plates as described above. hiPSC derived astrocytes were transduced with BFP lentivirus at plate-down at a MOI of 25 to demarcate from neurons. After 35 days astrocyte monocultures and after 21 days astrocyte-MN co-cultures were incubated with 100 nM of potentiometric dye MitoTracker Red CMXRos for 45 minutes. The cells were then washed with PBS and fixed in 4% PFA. Immunostaining was used to demarcate cell types with neuron-specific marker SMI312 and astrocyte specific marker GFAP (ab4674, Abcam). Fluorescence was visualised using Opera Phenix Plus high-content screening system at 40X magnification (OperaPHX/OPRTCLS Water Immersion Objective 40X: NA 1.1, WD 0.62 mm (field of view: approx. 323 μm x 323 μm). Flatfield correction was automatically performed by Harmony Software during image acquisition. Harmony was used for image analysis. Integrated fluorescent intensity of CMXRos in BFP+ cytoplasmic cell body area was calculated.

### Lentiviral overexpression of PGC1α

PGC1α overexpression lentivirus that was previously generated (6), was transduced in C9 hiPSC derived astrocytes at a multiplicity of infection (MOI) of 5.

### Statistical analysis

Statistical analysis for Figures 1-3 and Supplementary Figures 4 and 6 was performed using R version 4.3.2 (29). Generalised linear mixed models fit using the R package “lme4” (30), were employed. The random effects in GLMM define the nested structure of the data. This analytical approach is particularly suitable for addressing non-Gaussian data distributions and for accounting for the potential non-independence of measurements, which can lead to issues of pseudo-replication (31). We compared individual mutant iPSC lines to their corresponding gene-corrected isogenic controls, as illustrated in Figures 1-3 and Supplementary Figures 4 and 6. We also aggregated all measurements across the separate astrocyte lines to assess the significance of differences in astrocyte statuses (e.g., no astrocytes Vs. mutant astrocytes Vs. isogenic astrocytes), detailed in the Supplementary table 1. Throughout the latter analysis, we maintained the same considerations for non-independence as applied in the disaggregated analyses. In the GLMM model, post-hoc analyses were performed where appropriate to test for significance.

Statistical analysis for the remaining supplementary figures was performed using Prism version 8.4.0 (GraphPad Software). Data are presented as mean ± standard error of mean. Data were initially determined to be parametric or non-parametric before applying the appropriate statistical analysis, with false discovery rate (FDR) correction for multiple comparisons, as stated in the legends [\**p* < 0.05, \*\**p* < 0.01, \*\*\**p* < 0.001, \*\*\*\**p* < 0.001]. Throughout the study, independent biological replicates are defined as independently performed experiments.

## Acknowledgements

The authors thank following people providing technical assistance to this project: James Cooper, Karen Gladstone, Dr Arpan Mehta, Dr Alessandra Cardinali. Furthermore, we would like to thank Dr Pamela Brown and Linda Ferguson (Institute of Regeneration and Repair, University of Edinburgh) for generating lentiviruses.

## Author Contributions

MS, GEH, SC and BTS conceived and designed the study. MS, AH, RNC and ZJ performed data collection, analysis and interpretation. OD and MS performed bioinformatics and statistical analysis. KB, JN and DS provided technical cell-culture and laboratory expertise. JCW provided support with high-content imaging. GEH, SC, and BTS contributed to interpretation of data. MS drafted the manuscript, with critical input from GEH, SC, and BTS.

## Funding and Grants

MS is independently funded by the Medical Research Council (MRC) (MR/T000708/1) and the University of Edinburgh CMVM Institutional Bridging Funding for Clinical Academics. She also acknowledges support from the Rowling Scholars scheme, administered by the Anne Rowling Regenerative Neurology Clinic (ARRNC), University of Edinburgh, and a Seedcorn grant from the RS Macdonald Charitable Trust. The Hardingham, Chandran and Selvaraj laboratories are supported UK Dementia Research Institute (UK DRI-4001, UK DRI-4003 and UK DRI-4010 respectively), which receives its funding from UK DRI Ltd, principally funded by the MRC. SC also acknowledges funding from the ARRNC and My Name’5 Doddie Foundation. BTS is a Rowling-DRI fellow and further supported by MRC, The Humane Research trust.

## Competing interests’ statement

The authors have no declarations to make.

## Inclusion and Ethics Statement

All collaborators of this study have fulfilled the required criteria for authorship and have been included as authors, as their participation was essential for the design and implementation of the study. Roles and responsibilities were agreed among collaborators ahead of the research. This research was not severely restricted or prohibited in the setting of the researchers, and does not result in stigmatization, incrimination, discrimination or personal risk to participants. Research relevant to our study was considered in citations.

## Data availability statement

The datasets generated and analysed during the present study are available from the corresponding author upon request.

## Notes

### Competing Interest Statement

The authors have declared no competing interest.

