## Supplementary Figures 1-7 and Supplementary Table 1 for "Astrocytes control motor neuronal mitochondrial axonal transport deficits in C9ORF72 ALS"

Supplementary Figure 1

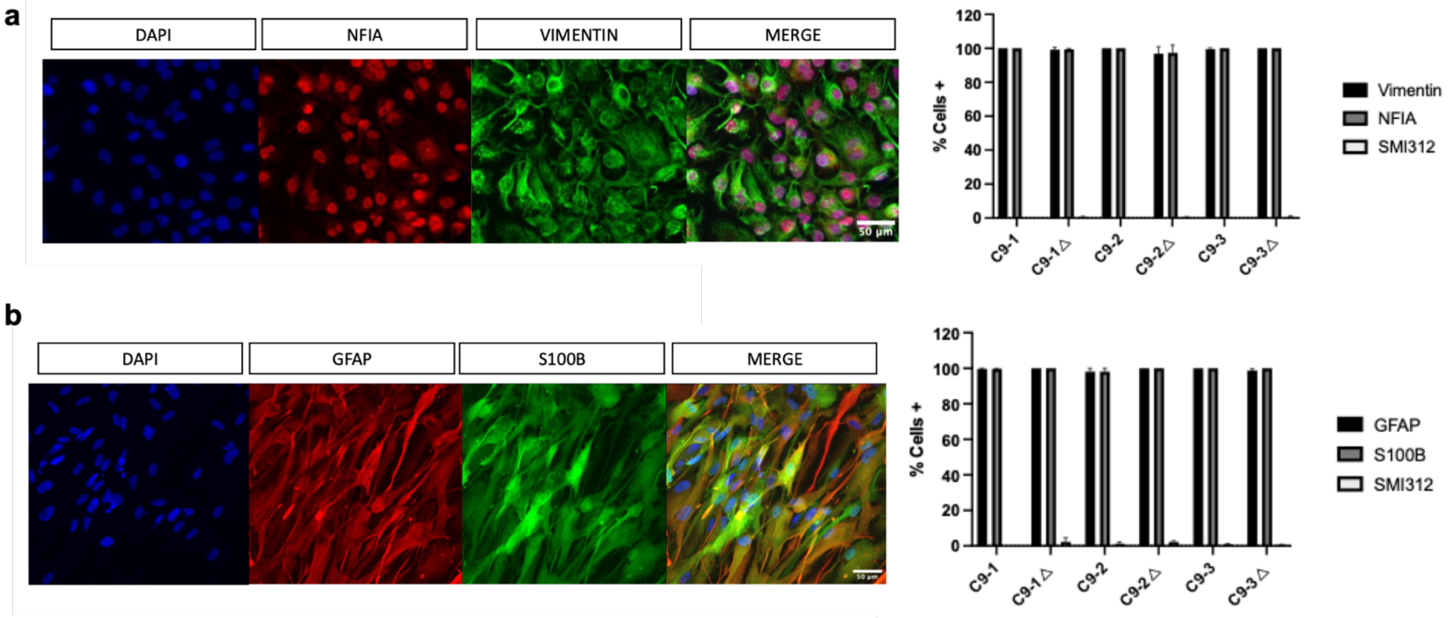


**Supplementary Figure 1:** Generation and characterisation of astrocyte progenitor cells (APCs) at 48-hours post-plating and of astrocytes, after two weeks of differentiation. Immunocytochemistry showed high expression of APC markers vimentin and nuclear factor I-A (NFIA), and quantitative immunolabeling at 2 weeks post-differentiation revealed >90% of cells positive for astrocyte markers, S100 calcium-binding protein B (S100B) and glial fibrillary acidic protein (GFAP), alongside <5% minimal expression of SMI312, a neuronal marker. **(a)** Representative images of vimentin and NFIA immunostaining in astrocyte progenitors (upper panel) and S100B and GFAP immunostaining in 2-week-old astrocyte cultures (lower panel); (Scale bars: 50 µm). **(b)** Percentage of S100B$^+$, GFAP$^+$ and SMI312$^+$ cells in 2-week-old astrocyte cultures derived from the three pairs of iPSC lines ($N$ = 3–7, at least 400 cells per cell line per experiment, one-way ANOVA with Bonferroni correction).

Supplementary Figure 2

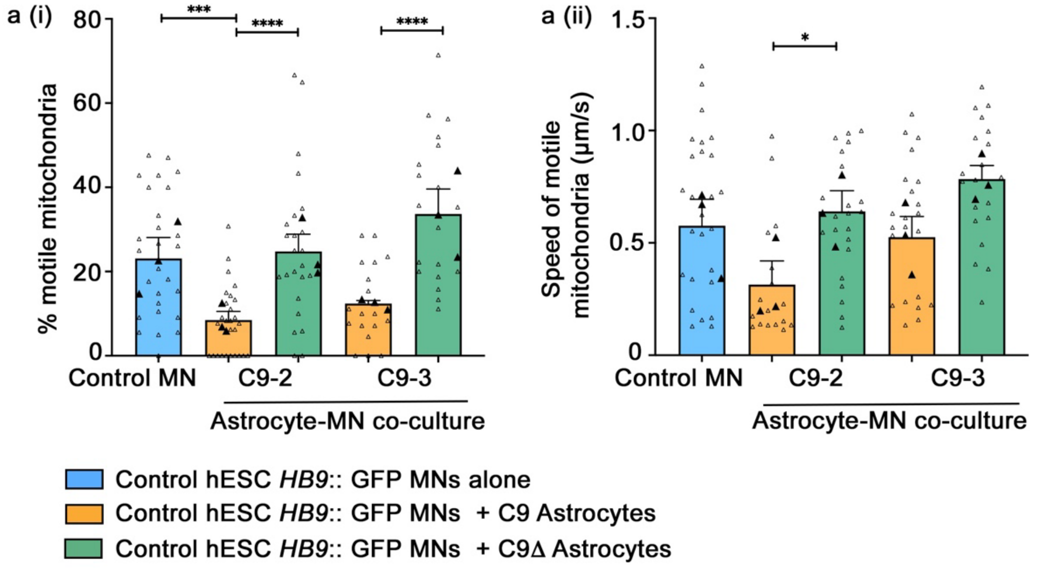


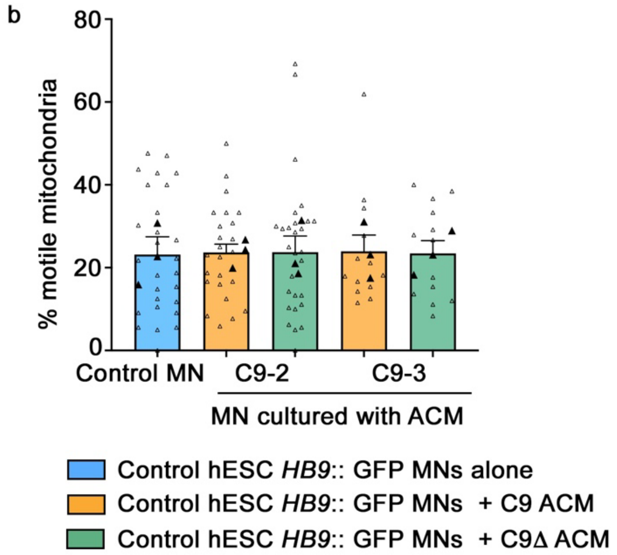

**Supplementary Figure 2. (a)** Effects of astrocytes on mitochondrial transport in control hESC *HB9*::GFP MNs in physical co-cultures. **(i)** Quantification of the percentage of motile mitochondria (labelled with mitoDsRed2) and **(ii)** mean speed of motile mitochondria in a 100 µm stretch of proximal axon in control hESC *HB9*::GFP MNs co-cultured with either C9-astrocytes or gene-corrected astrocyte controls (C9Δ-A) from two independent patient lines. The open triangles represent technical replicates (individual axons) and the filled triangles represent independent biological replicates (mean of technical replicates). Data are represented as the mean of biological replicates ± SEM of the means of biological replicates**.** p-values (*<0.05; **<0.01; ***<0.001; ****<0.0001) determined by fitting generalised linear mixed models, followed by post-hoc tests with Bonferroni correction. **(b)** Quantification of the percentage of motile mitochondria (labelled with mitoDsRed2) in a 100 µm stretch of proximal axon in control hESC *HB9*::GFP MNs treated with conditioned medium from either C9-astrocytes or C9Δ-A from two independent patient lines. The open triangles represent technical replicates (individual axons) and the filled triangles represent independent biological replicates (mean of technical replicates). Data are represented as the mean of biological replicates ± SEM of the means of biological replicates**.** Data are analysed by fitting generalised linear mixed models, followed by post-hoc tests with Bonferroni correction.

Supplementary Figure 3

a

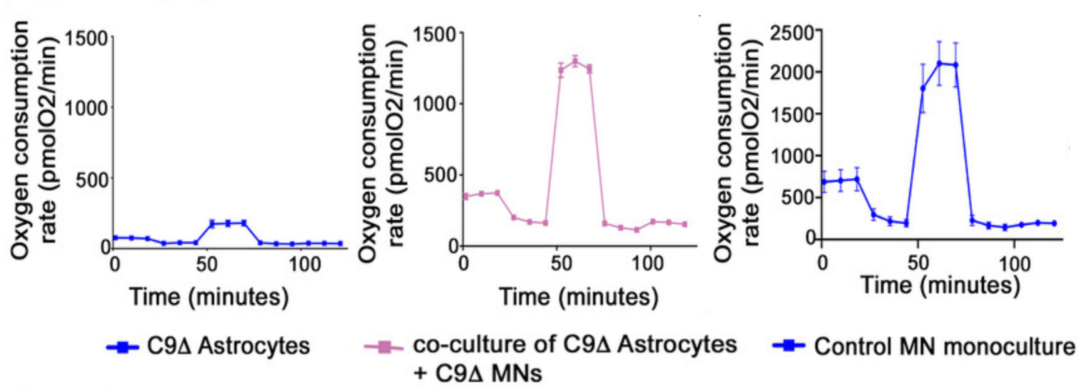


b

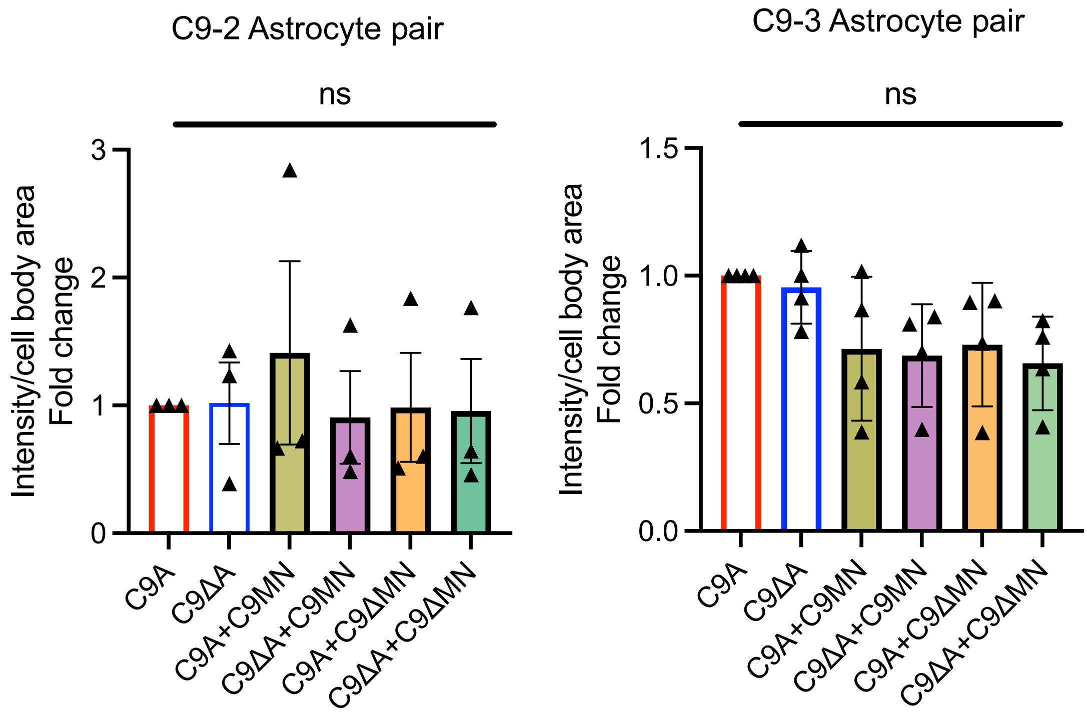

**Supplementary figure 3. (a)** Evaluating the feasibility of using the seahorse assay to assess how C9 astrocytes affect mitochondrial respiratory activity. The mitochondrial OCR of MN monocultures and MNs co-cultured with astrocytes were nearly 8-fold higher than astrocytes alone (plated cell densities for: astrocytes – 200,000 cells; MNs – 200,000 cells; astrocyte-neuronal co-cultures – 200,000 cells of each cell type). **(b)** Quantification of the mean fluorescence intensity of MitoTracker Red CMXROS in hiPSC-derived astrocytes (from the three-independent patient-derived C9orf72 lines) in monocultures and in co-cultures with hiPSC-derived MNs (one C9-mutant/gene-corrected pair). This figure summarises data from three or more independent biological experiments (plate downs) [N ≥ 3] with each symbol representing the average from n = 2-4 wells per condition per plate down. Data are represented as mean ± SEM. Normality testing indicated that the data follows a normal distribution. Consequently, a one-way ANOVA was conducted to assess differences among group means.

Supplementary Figure 4

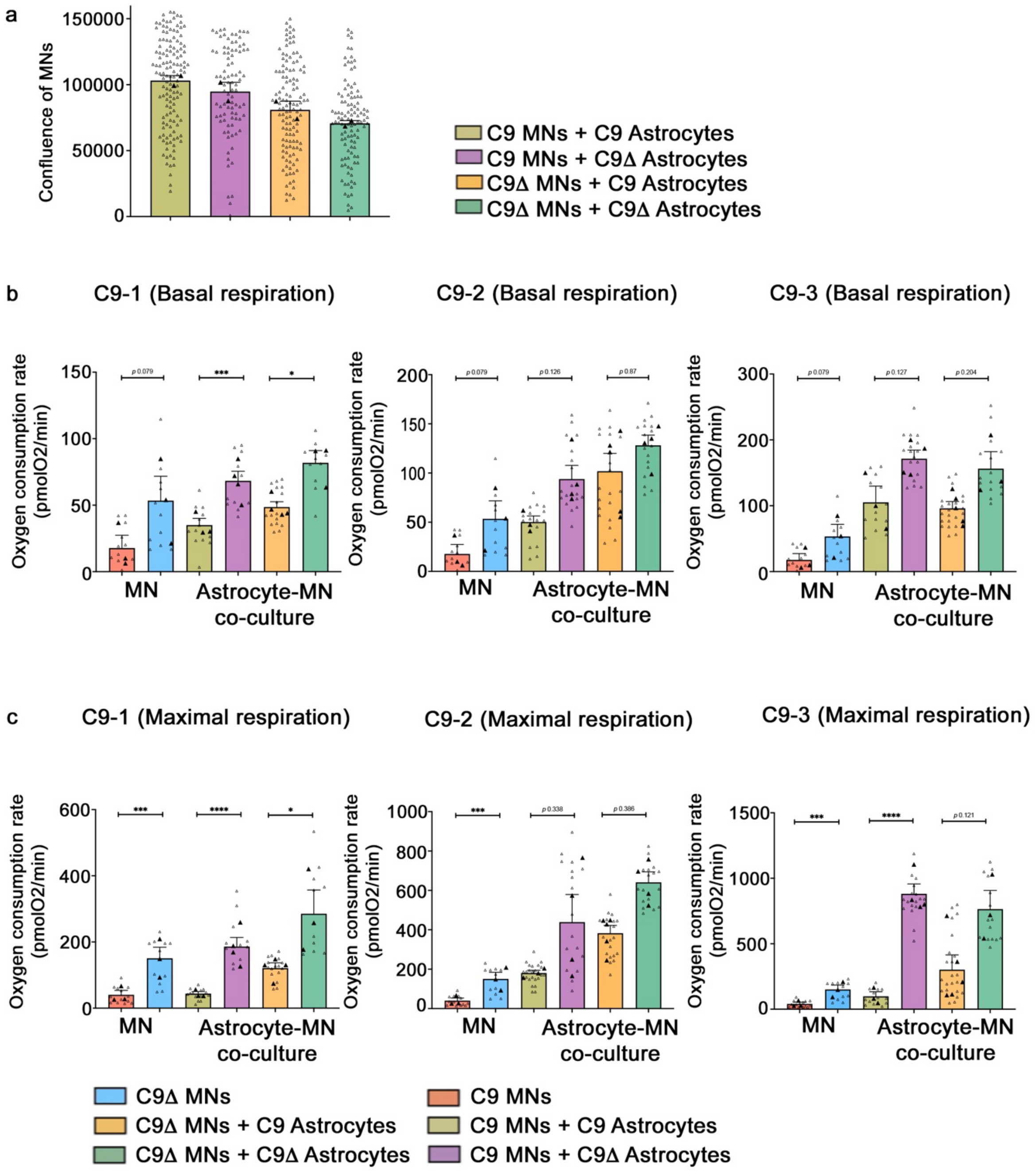

**Supplementary figure 4. (a)** Neuronal confluence in co-culture experiments was determined by neuron-specific staining via SMI-312 immunocytochemistry. **(b) & (c)** Co-cultures with astrocytes, regardless of the genotype, enhanced basal and maximal respiration compared to MN monocultures. Plated cell densities for: MNs – 50,000 cells; astrocyte-neuronal co-cultures – 50,000 cells of each cell type. The open triangles represent technical replicates, and the filled triangles represent independent biological replicates (mean of technical replicates). Data are represented as the mean of biological replicates ± SEM of the means of biological replicates**.** p-values (*<0.05; **<0.01; ***<0.001; ****<0.0001) determined by fitting generalised linear mixed models, followed by post-hoc tests with Bonferroni correction.

Supplementary Figure 5

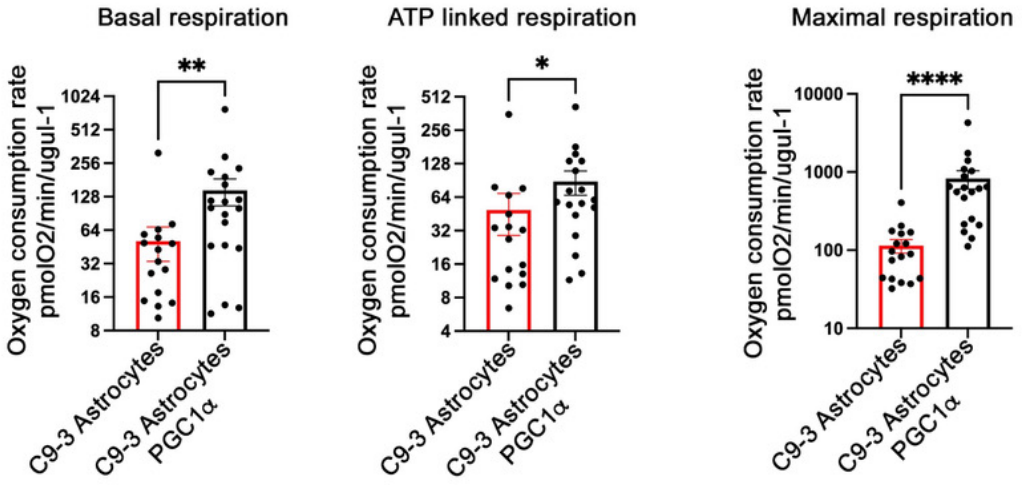


**Supplementary Figure 5.** Boosting astrocyte metabolism with selective PGC1α pathway manipulation in C9 astrocytes increases basal, ATP-linked and maximal respiration. Data are represented as mean ± SEM; n ≥ 3 wells per line per experiment, with experiments repeated in three independent cultures from different differentiations (N = 3). P-values (*<0.05; **<0.01; ***<0.001; ****<0.0001) determined by Mann Whitney U non-parametric test.

Supplementary Figure 6

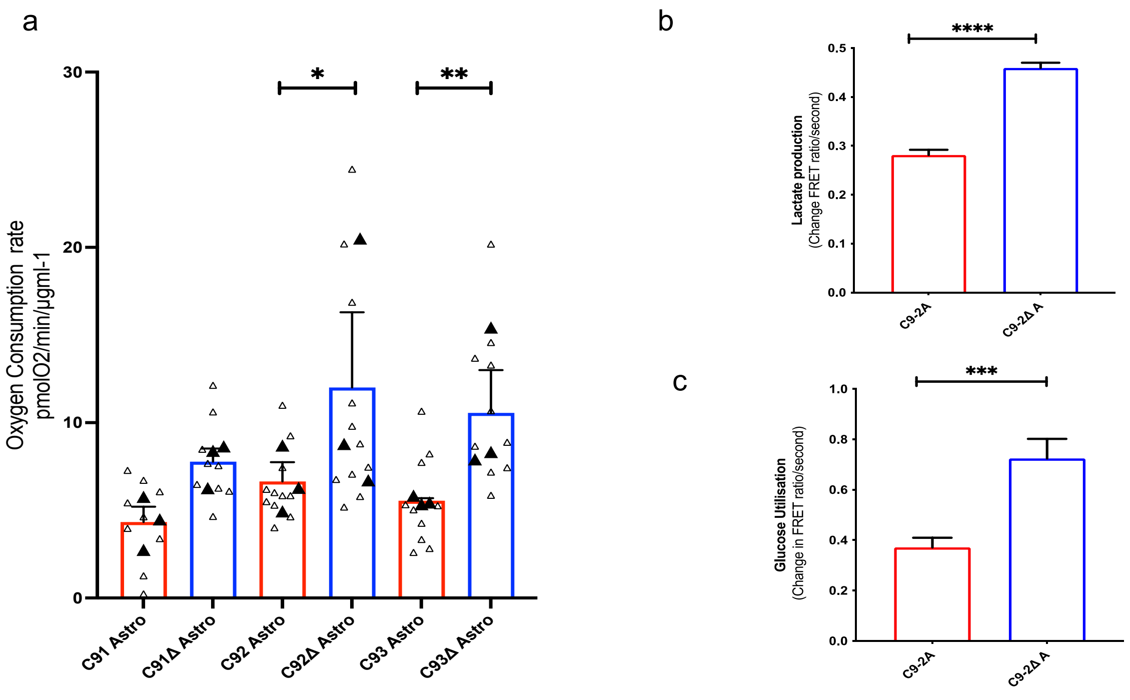


**Supplementary Figure 6:** C9 astrocytes, when compared to their corresponding gene-edited controls, display **(a)** glycolytic impairment and, **(b)** diminished lactate production and **(c)** glucose utilisation. The open triangles represent technical replicates (wells studied per plate down) and the filled triangles represent independent biological replicates (mean of technical replicates). Data are represented as the mean of biological replicates ± SEM of the means of biological replicates**.** For (a) P-values (*<0.05; **<0.01; ***<0.001; ****<0.0001) are determined by one-way ANOVA following normality testing indicating that the data follows a normal distribution. For (b) and (c) P-values (*<0.05; **<0.01; ***<0.001; ****<0.0001) are determined by unpaired t-test.

Supplementary Figure 7

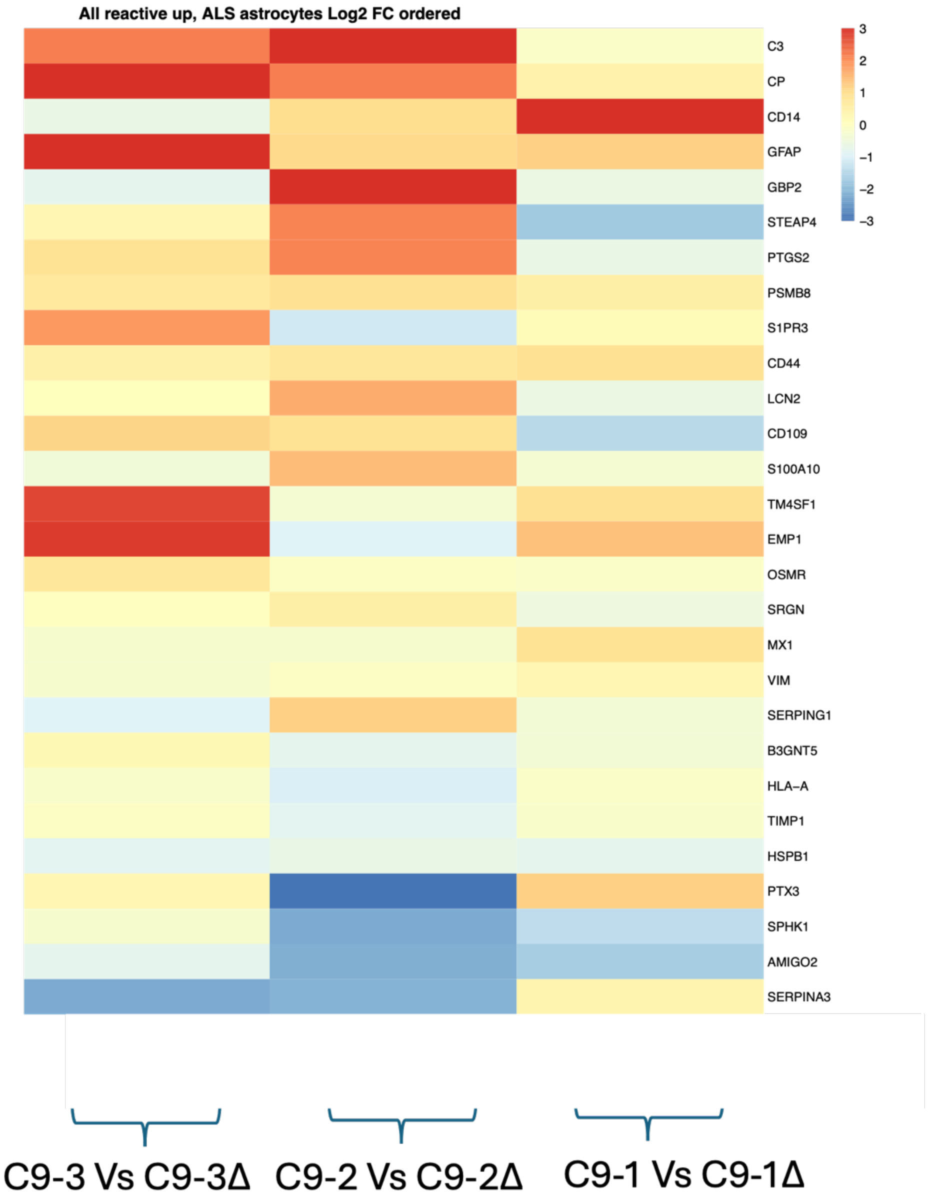


**Supplementary Figure 7.** Analysis of RNA sequencing on hiPSC-derived astrocytes from the three independent C9ORF72 mutant lines and their isogenic gene-corrected controls to determine genotype-driven astrocyte reactivity using gene sets for astrocyte reactivity from Zhang et al. (2014) (32). Our analysis revealed no consistent effect of the C9 mutation on astrocyte reactivity at the experimental time point (14 days post-differentiation). Log2 fold-change (log2FC) heatmaps are generated using DESeq2 (33) to compare mutant and isogenic gene-corrected samples for each gene. In the heatmap, blue indicates lower expression in the mutant line compared to isogenic gene-corrected control, red indicates higher expression in the mutant line, and yellow represents similar expression between mutant and isogenic control. Log2 fold-change values have been constrained to a range of -3 to +3 for visualisation. Additionally, purple/grey indicates where DESeq2 could not calculate a fold change.

Supplementary table 1

Combined Analysis

**Fig 1c:**

| Comparison | Standard Error (SE) | z-value | p-value | Significance |
|---|---|---|---|---|
| Control MNs – Co_ast_mut | 0.36138 | 3.20 | 0.0041 | *** |
| Control MNs – Co_ast_iso | 0.36500 | -0.24 | 1.0000 | Not significant |
| Co_ast_mut – Co_ast_iso | 0.24484 | -5.08 | <0.0001 | *** |

**Fig 1d:**

| Comparison | Standard Error (SE) | z-value | p-value | Significance |
|---|---|---|---|---|
| Mutant MNs – Co_ast_mut | 0.2885 | -0.76 | 1.0 | Not significant |
| Mutant MNs – Co_ast_mut | 0.2947 | -4.83 | 4.1e-06 | *** |
| Co_ast_mut – Co_ast_iso | 0.2030 | -5.93 | <0.0001 | *** |

**Fig 1e:**

| Comparison | Standard Error (SE) | z-value | p-value | Significance |
|---|---|---|---|---|
| Control MNs - Co_ast_mut | 0.2464 | 3.01 | 0.0079 | ** |
| Control MNs - Co_ast_mut | 0.2511 | -0.53 | 1.0000 | Not significant |
| Co_ast_mut – Co_ast_iso | 0.1664 | -5.25 | <0.0001 | *** |

**Fig 1f:**

| Comparison | Standard Error (SE) | z-ratio | p-value | Significance |
|---|---|---|---|---|
| Mutant MNs – Co_ast_mut | 0.245 | 3.795 | 0.0004 | *** |
| Mutant MNs – Co_ast_mut | 0.251 | -0.430 | 1.0000 | NS |
| Co_ast_mut – Co_ast_iso | 0.177 | -5.881 | <0.0001 | *** |

**Fig 2a:**

| Contrast | SE | z.ratio | p.value |
|---|---|---|---|
| ast_mut.mn_mut - ast_iso.mn_mut | 0.1612224 | -3.7061104 | 0.0008419 |
| ast_mut.mn_mut - ast_mut.mn_iso | 0.1563660 | -1.9761775 | 0.1925387 |
| ast_iso.mn_mut - ast_iso.mn_iso | 0.1564306 | -0.9361951 | 1.0000000 |
| ast_mut.mn_iso - ast_iso.mn_iso | 0.1513950 | -2.8729536 | 0.0162662 |

**Fig 2b:**

| Contrast | SE | z.ratio | p.value |
|---|---|---|---|
| ast_mut.mn_mut - ast_iso.mn_mut | 0.2643360 | -5.426509 | 0.0000002 |
| ast_mut.mn_mut - ast_mut.mn_iso | 0.2655309 | -3.807911 | 0.0005606 |
| ast_iso.mn_mut - ast_iso.mn_iso | 0.2508583 | -1.221491 | 0.8876013 |
| ast_mut.mn_iso - ast_iso.mn_iso | 0.2510797 | -2.906349 | 0.0146269 |

**Fig 2d:**

| Contrast | SE | z.ratio | p.value |
|---|---|---|---|
| ast_mut.mn_mut - ast_iso.mn_mut | 0.2558018 | -5.0091359 | 2.2e-06 |
| ast_mut.mn_mut - ast_mut.mn_iso | 0.2460399 | 0.1831919 | 1.0e+00 |
| ast_iso.mn_mut - ast_iso.mn_iso | 0.2562191 | -0.2552255 | 1.0e+00 |
| ast_mut.mn_iso - ast_iso.mn_iso | 0.2457843 | -5.6627393 | 1.0e-07 |

**Fig 2e:**

**Basal Respiration**

| Effect | Chi-squared | Degrees of Freedom | p-value |
|---|---|---|---|
| astro_status | 17.558 | 1 | 2.787e-05 |

**ATP-linked Respiration**

| Effect | Chi-squared | Degrees of Freedom | p-value |
|---|---|---|---|
| astro_status | 20.238 | 1 | 6.839e-06 |

**Maximal Respiration**

| Effect | Chi-squared | Degrees of Freedom | p-value |
|---|---|---|---|
| astro_status | 9.4659 | 1 | 0.002093 |

**Fig 3b:**

| Comparison | Standard Error (SE) | z-value | p-value | Significance |
|---|---|---|---|---|
| Control MNs – Co_ast_mut | 0.2314 | 4.98 | 1.9e-06 | *** |
| Control MNs – Co_ast_pgc1a | 0.2325 | -0.88 | 1.0 | Not significant |
| Co_ast_mut – Co_ast_pgc1a | 0.1633 | -8.31 | <0.0001 | *** |

**Fig 3c:**

| Comparison | Standard Error (SE) | z-value | p-value | Significance |
|---|---|---|---|---|
| Mutant MNs – Co_ast_mut | 0.2318 | -1.06 | 0.8709 | Not significant |
| Mutant MNs – Co_ast_pgc1a | 0.2327 | -6.24 | <0.0001 | *** |
| Co_ast_mut – Co_ast_pgc1a | 0.1578 | -7.65 | <0.0001 | *** |

**Fig 3d:**

| Comparison | Standard Error (SE) | z-value | p-value | Significance |
|---|---|---|---|---|
| Control MNs – Co_ast_mut | 0.2875 | 2.61 | 0.0244 | ** |
| Control MNs – Co_ast_pgc1a | 0.2939 | -0.94 | 0.615 | Not significant |
| Co_ast_mut – Co_ast_pgc1a | 0.1892 | -5.43 | <0.0001 | *** |

**Fig 3e:**

| **Comparison** | **Standard Error (SE)** | **z-value** | **p-value** | **Significance** |
|---|---|---|---|---|
| Mutant MNs – Co_ast_mut | 0.2560 | 3.69 | 0.0007 | *** |
| Mutant MNs – Co_ast_pgc1a | 0.2647 | -1.39 | 0.348 | Not significant |
| Co_ast_mut – Co_ast_pgc1a | 0.1965 | -6.67 | <0.0001 | *** |

**Significance Codes:**

- ***: p < 0.001
- **: 0.001 < p < 0.01
- *: 0.01 < p < 0.05
